# Quantum Magnetization Exchange through Transient Hydrogen Bond Matrix Defines Magnetic Resonance Signal Relaxation and Anisotropy in Central Nervous System

**DOI:** 10.1101/2025.04.22.650086

**Authors:** Dmitriy A. Yablonskiy, Alexander L. Sukstanskii

**Affiliations:** Mallinckrodt Institute of Radiology, Washington University in St. Louis, 4525 Scott Ave. Room 3216, St. Louis MO, 63110; Hope Center for Neurological Disorder, 660 S. Euclid Ave., St. Louis, Missouri 63110; Knight Alzheimer Disease Research Center, 4488 Forest Park Ave., St. Louis, MO 63108; Department of Biomedical Engineering, Washington University in St. Louis, 1 Brookings Drive, St. Louis, MO 63130

**Keywords:** MRI, Transient Hydrogen Bonds, Myelin, MR signal relaxation, Cellular membranes, Neurons

## Abstract

The integrity of cellular membranes (lipid bilayers) and myelin sheaths covering axons is a crucial feature controlling normal brain structural and functional networks. Yet, in vivo evaluation of this integrity at the nanoscale level of the cellular membranes organization is challenging.

Herein we explore the dual property of biological water in Central Nervous System (CNS), as one of the major stabilizing factors of cellular membranes, and the major source of MRI signal. We introduce the Basic Transient Hydrogen Bond (THB) model of the MR signal relaxation due to the quantum spin/magnetization exchanges within the THB Matrix encompassing water molecules and membrane-forming macromolecules.

Our data show the existence of two THB Matrix components with distinct lifetimes – one in a few nano-second range, and another in the range of tens nanoseconds. Importantly, the former component facilitates longitudinal relaxation of MR signal, the latter contributes to its transverse relaxation and causes the anisotropy of MR signal relaxation. These distinct features offer opportunity to study nanoscale level microstructure of cellular membranes. Furthermore, the ability to differentiate distinct THB Matrix components based on their MR signal relaxation properties can be fundamental to identifying pathological changes and enhancing disease visibility on MRI scans.

## INTRODUCTION

Since introduction by Lauterbur [1], MRI plays an increasingly profound role in studying biological tissue properties in humans and animals in health and disease. Among numerous MRI techniques developed to highlight different properties of biological tissue microstructure and functioning, images based on relaxation tissue properties (longitudinal - T1 and transverse - T2) most often serve in clinical practice for highlighting biological tissue contrast in different human (and animal) organs in healthy and clinically pathological cases. Clearly, understanding biophysical mechanisms underlying water molecules (the major source of MRI signal) interaction with the elements of tissue cellular structure is required for deciphering biological meaning of MRI images.

Quite a few models have been developed over the years to elucidate relationships between tissue microstructure and MR signal relaxation properties in neuronal tissue. Apparently, to describe biophysical sources of MR signal relaxation we need to establish a mechanism of magnetization exchange between water protons (aqueous component of biological tissue) and protons forming tissue cellular composition (often called semisolid, or bound protons). Such an exchange can take place either by means of physical exchange of protons (hence, their spin magnetizations), or by means of the protons’ spin exchange due to their quantum spin-spin interactions. While the former process is usually described in terms of the Bloch-McConnell equations, the latter process can be described in the framework of the theory of quantum spin exchanges between proton dipoles proposed by Bloembergen, Purcell, and Pound [2]. The majority of existing models so far rely on the Bloch-McConnell equations introduced by Morrison et al [3] to describe Magnetization Transfer (MT) experiments. The Bloch-McConnell equations result from merging the Bloch equations describing MR signal relaxation dynamics in a single-component media, with the particle exchange mechanism proposed by McConnell [4] for treating chemical reactions. By providing a convenient framework for analyzing experimental data with different level of sophistication [3, 5-16], these models do not explain the biophysical mechanisms defining phenomenological relaxation and cross-relaxation models’ parameters. Accordingly, such models cannot explain experimentally-established magnetic field dependence of MR signal relaxation [17-19] and its anisotropic features [20-33] in neuronal tissues.

A natural approach to developing theory of MR signal relaxation would rely on magnetization exchange between aqueous and bound protons by means of quantum dipole interactions, which are highly sensitive to magnetic field strength and the relative orientations of the interacting dipoles [2]. Given the strong distance dependence of the dipole-dipole interactions, such a theory would necessitate a detailed understanding of tissue microstructure at the nanoscale level. Indeed, while molecular dynamic simulations within the myelin sheath [34] have highlighted the importance of actual tissue structure in determining MR signal decay, the recently introduced Transient Hydrogen Bond (THB) model [35] offers a significant advantage by providing a more analytically tractable framework. This model yielded analytical equations that accurately described T1 signal evolution, offering a valuable tool for understanding and predicting MR signal relaxation behavior.

The THB model [35] relies on specific information relating neuronal tissue cellular structure to its magnetic properties. In early publications, Koenig et al [36, 37] suggested cholesterol as an important player in formation of gray-white matter contrast of MRI signal. Kucharczyk et al [38] emphasized that the phosphatidylcholine component of myelin can play even more important role on relaxation of MRI signal. This was in agreement with results of Ceckler et al [39] who demonstrated by experiments with bilayers of sphingomyelin (one hydroxyl group), phosphotidylglycerol (two hydroxyl groups), and phosphotidylinositol (five hydroxyl groups) that the magnitude of the MT effect increased linearly with the number of hydroxyl groups per unit area. The studies by Bryant and Korb [40] and Halle [41] emphasized the role of magnetic coupling of protons with motion-immobilized species on T1 relaxation.

Based on these findings, the THB model [35] attributes the major features of MRI signal relaxation properties in brain tissue to organization of lipid bilayers in myelin sheath and cellular membranes. Specifically, the THB model considers hydrogen bonding of water molecules transiently trapped in hydrophilic heads of cellular membranes and myelin-forming lipid bilayers as a biophysical mechanism for the dipole-dipole interaction between *sub-populations of bound protons in the hydrophilic heads* and *water protons* from adjacent to membranes spaces, i.e., intracellular, extracellular and myelin water sub-layers. Hence, the term Transient Hydrogen Bond (THB) model. It is important to emphasize that only macromolecular-bound protons available for transient hydrogen bonding with water protons (such as residing in hydrophilic heads) contribute to the spin-exchange processes, while not-available bound protons (such as locating in the hydrocarbon cores) do not. The microscopic theory of MR signal relaxation of the lipid-bound protons due to the lateral diffusion of lipid chains, the Lateral Diffusion Model (LDM), was developed previously [42].

In present paper, we focus on the role of quantum dipole interactions in formation of T1 and T2 contrast in brain neuronal tissue in the framework of the THB model [35] accounting for relaxation effects of biological tissue aqueous water proton magnetization by means of their quantum dipole interactions with bound protons of cell-building macromolecules. These interactions take place when water molecules, in the process of their motion through the biological tissue (by means of diffusion or facilitated transfer through the aquaporins in the cellular membranes), form transient hydrogen bonds with macromolecules. The THB model has two major parameters – *λ* (the strength of quantum interactions within a THB, Eq. (31)) and *τ* (the THB lifetime). It turns out that the contribution of THBs with short *τ* (a few nanoseconds that we reported in [35]) describes the T1 field dependence pretty accurate but provides small contribution to the T2 signal relaxation. Herein we introduce two types of THBs – with short and long *τ* (the latter has a small contribution to T1 but a quite significant contribution to T2). We validate our new model using previously published data [43].

In the current paper, we consider only the quantum dipole-dipole interactions and their contribution to T1 and T2 relaxation. There are other important contributions (mainly, to T2 and T2*), especially at high *B*_0_ fields, related to the presence of iron in tissue (see [44, 45] and follow-up publications); however, this effect is beyond the scope of the current manuscript.

## THEORETICAL BACKGROUND

### Biophysical underpinnings of the Transient Hydrogen Bond (THB) model

Cellular membranes are the major parts of biological tissue structure. They separate intracellular machinery from extracellular fluid encompassing all cells and cellular processes. While the central part of lipid bilayers is hydrophobic and the water concentration in this part of the membrane is very low (about 10^−4^ of its concentration in the surrounding space), the external parts of membranes, i.e., lipid heads, are hydrophilic. Quite a few papers using molecular dynamic approach demonstrated that about 25% of the cellular membrane’s outer part is occupied simultaneously by protons bound to macromolecules and water protons [46, 47]. This offers the favorable environment for interaction between head-bound protons and free water protons. Importantly, the polar sugar ring headgroups of GC and other glycolipids are extended outward from the membrane surface [48], thus effectively increasing the thickness of the interface region and permitting stronger interaction of the galactose headgroup hydroxyls with bulk water molecules. This interaction leads to formation of transient hydrogen bonds between water and corresponding counterparts of bilayers (oxygen, nitrogen, and protons). In cellular membranes, more than 50% of galactosylceramide and sulfatide are hydroxylated at the 2-C atom by fatty acid 2-hydroxylase (FA2H), thereby providing additional hydroxyl groups for hydrogen bonding [49, 50].

### THB Model Master Equations

Let’s consider a general case with several types of transient hydrogen bonds (THB) with different bond lifetimes *τ* _*j*_, *j* = 1, 2, …, *N*_*TR*_, where *N*_*TR*_ is the total number of THB types. Let *N*_*bj*_ denote the total number of bound protons of type *j* in lipid heads (per unit volume), with fraction *n*_*bj*_ participating in the *dipole-dipole interaction with water protons* at any given moment. Let *N*_*w*_ denote the total number of water protons (per unit volume), with a fraction *n*_*wj*_ in the *j*-type THB at any given moment. The following relationships describe the balances of interacting protons for each THB type:

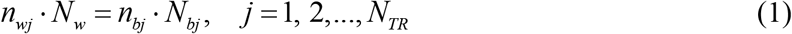

As already mentioned above, only bound protons in lipid bilayers residing in hydrophilic lipid heads can come in close contact with water protons from hydrogen-bound water molecules. The bound protons in the vicinity of hydrophobic lipid tails have minor contribution to the water-lipid interaction because water concentration in the lipid core is several orders of magnitude smaller than in and around lipid tails.

According to the above consideration of interacting pools, the THB theory [35] can be generalized to the two sets of systems of *N*_*TR*_ +1 cross-connected equations for the longitudinal (*z*, Eqs. (2)) and transverse (⊥, Eqs. (3)) magnetizations:

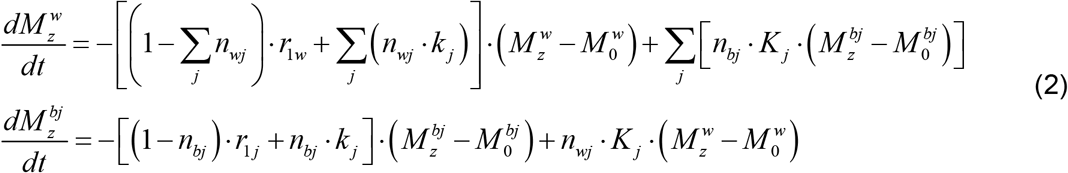

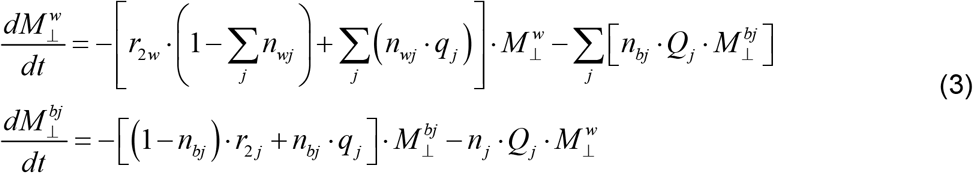

In each of the systems, one equation corresponds to the water protons (index *w*) and *N*_*TR*_ equations (indices *bj*) correspond to *b*-protons in the *j*^th^ type of THB. The sums are taken over all THBs, 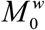 and 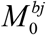 are equilibrium magnetizations of water and *j*-components of bound protons, *r*_1*w*_, *r*_1 *j*_ *r*_2*w*_ and *r*_2*j*_ are longitudinal and transverse relaxation rates defined by interactions other than participating in the *w-b* THBs. The diagonal, *k*_*j*_, *q*_*j*_, and cross-relaxation, *K* _*j*_, *Q*_*j*_, coefficients are defined by Eqs. (28) and (29) in the Appendix 1.

Note a different sign of the exchange terms *Q*_*j*_ in Eq. (3) as compared to the exchange terms *K* _*j*_ in Eq. (2). Also note a different correlation functions’ dependence of the diagonal and cross-exchange terms in Eqs. (29) and (28).

In deriving Eqs. (2) and (3), we considered that the water protons are mobile and can interact with plurality of bound protons during the course of MR signal evolution, while the bound protons are components of the long-chain lipids and proteins, their mobility is significantly restricted, so those bound protons can interact only with bound protons of the same type.

Given the complexity of an exact solution to Eqs. (2) and (3), we employ perturbation theory, leveraging the small magnitude of the parameters *n*_*wj*_ (concentrations of THBs). In the first approximation with respect to *n*_*wj*_ ≪ 1, a solution to Eqs. (2) for water longitudinal magnetization is as follows:

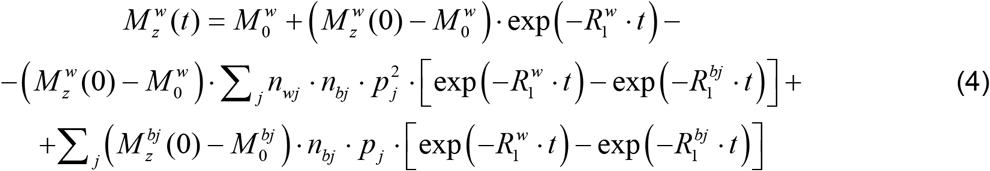

A solution for bound protons is:

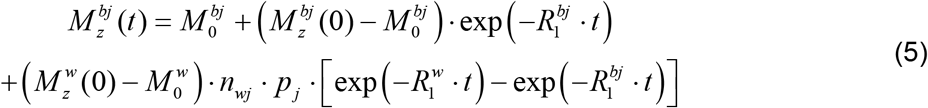

where 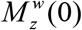 and 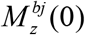 are the initial after the RF pulse magnetizations of water and *j*-type bound protons. The relaxation rate parameters for water, 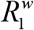, and bound protons, 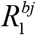 are:

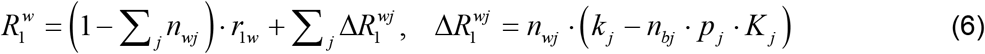

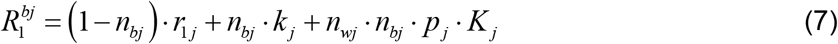

where the diagonal, *k* _*j*_, and cross-relaxation, *K* _*j*_, coefficients are defined by Eqs. (28) in the Appendix 1, and

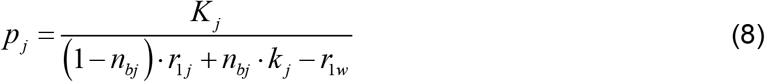

The solution for the transverse magnetization is as follows:

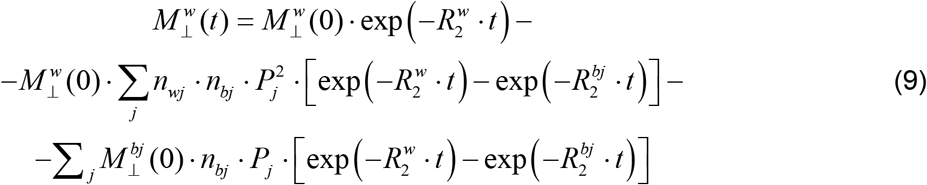

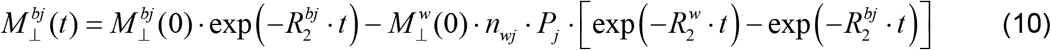

where 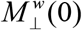 and 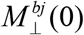 are the initial (after the RF excitation pulse) transverse magnetizations of water and *j*-components of bound protons. The relaxation rate parameters for water, 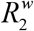, and bound protons, 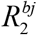, are:

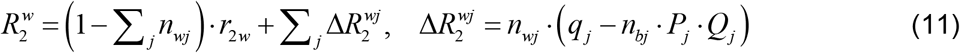

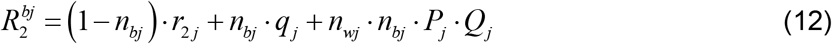

where the diagonal, *q* _*j*_, and cross-relaxation, *Q*_*j*_, coefficients are defined by Eqs. (29) in the Appendix 1, and

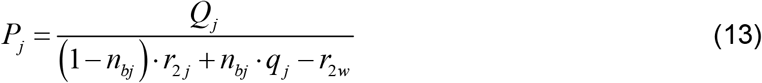

The above equations describe a general case valid for an arbitrary number of bound proton types. To address the major features of T1 and T2 relaxation, in what follows, we will consider a basic model that includes only two types of bound protons.

## RESULTS

### Basic THB Model of the Longitudinal (T1) Relaxation in CNS

As was demonstrated in [35], explaining anisotropic properties of T1 signal relaxation required the presence of THBs with structural orientation order in the THB dipole-dipole connections. At the same time, the anisotropy of T1 relaxation is rather small, hence the model cannot be restricted to a single anisotropic *b*-proton pool, but should include both, isotropic and anisotropic *b*-pools. In our approach, we characterize *b-*protons by two effective magnetizations: 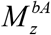 - anisotropic (*A*) pool comprising THBs forming orientation order with the dipole-dipole *w-b* connections oriented perpendicular to the surfaces of bi-layers surrounding axons, and 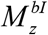 isotropic (*I*) pool with a random (uniform) distribution of the dipole-dipole *w-b* connections. The latter can originate from randomly oriented dipole-dipole connections in the axons and/or randomly oriented cellular membranes.

According to the above considerations, the basic model is described by the Eqs. (4)-(8) with index *j* taking only two values, *j* = *I, A*. Hence, for the total longitudinal magnetization the result is:

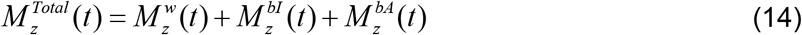

where each of the terms in Eq. (14) is proportional to the corresponding number of spins (*N*_*w*_, *N*_*bI*_, *N*_*bA*_) satisfying the relationships:

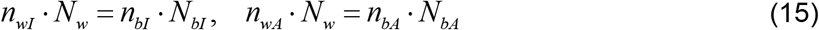

The diagonal and cross-relaxation coefficients are detailed in the Appendix 2. Note that despite seeming complexity, all the signals depend only on three exponentials: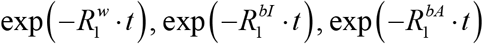.

The *isotropic contribution* to the MR signal is defined by the isotropic relaxation parameters:

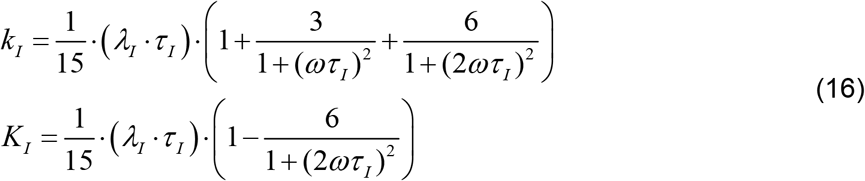

The contribution of isotropic THBs to 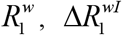, and the dependence of relaxation parameter 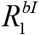 on the lifetime *τ* _*I*_ and on the model parameters are illustrated in Figure 1 and Figure 2 for different values of the external magnetic field *B*_0_.

**Figure 1.**
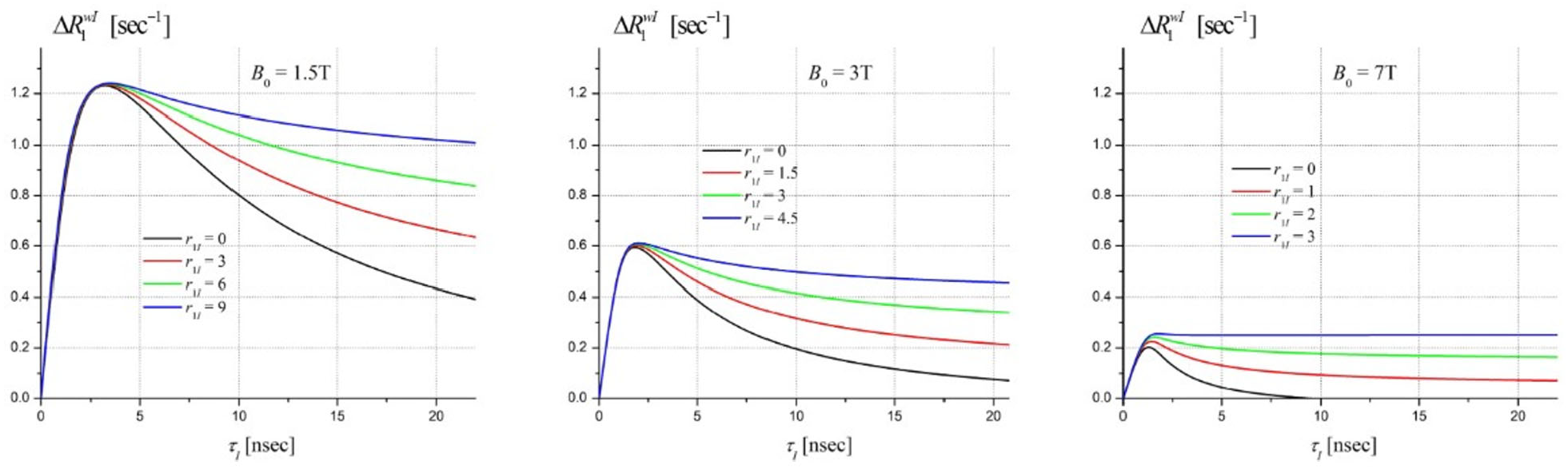
The contribution of the isotropic term 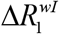 in Eq. (6) to 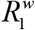 for different magnetic fields *B*_0_ as a function of the lifetime *τ*_*I*_ for *λ* = 20 (sec· nsec)^−1^, *n* _*wI*_ = 0.1, *n* _*bI*_ = 0.5, *r*_1*w*_ = 0.24 sec^−1^ and different values of the parameter *r*_1*I*_ (in sec^−1^, shown by the numbers on the plots).

**Figure 2.**
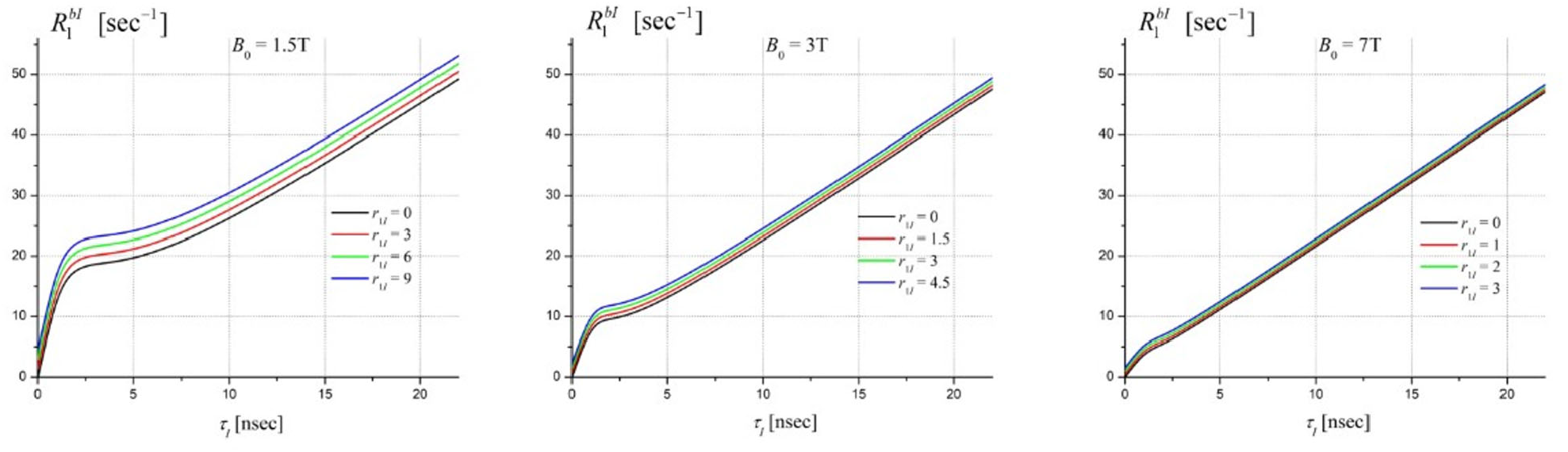
The dependencies of the isotropic relaxation parameter 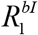 (Eq. (7) with *j=I*) on model parameters for different external magnetic fields *B*_0_. Plots illustrate examples of these dependences on the lifetime *τ* for *λ* = 20 (sec· nsec)^−1^, *n* _*wI*_ = 0.1, *n* _*bI*_ = 0.5, *r*_1*w*_ = 0.24 sec^−1^.

According to Eq. (16), the cross-relaxation parameter *K*_*I*_ < 0 for 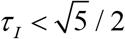 and *K*_*I*_ > 0 for 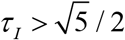. Hence, the terms proportional to *p*_*I*_ in Eqs. (6) and (7) are small for *ω* ·*τ* _*I*_ ≤ 2, and the *τ*-dependences of 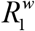and 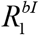 in this interval are defined mostly by the diagonal relaxation parameter *k*_*I*_. This explains why the isotropic contribution to 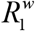 in Figure 1 is practically independent of *r*_1*I*_ in this interval and why the dependence of 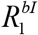 on *r*_1*I*_ is mainly described by the contribution of the additive term (1− *n*_*bI*_)· *r*_1*I*_.

*The anisotropic behavior* of relaxation parameters is determined by the combined anisotropy of the THB model and the relaxation parameter *r*_1 *A*_. The former is described by Eqs. (6) and (7) with notations defined in Appendix 2. There, Eqs. (33) define the general expressions for diagonal and cross-relaxation parameters, while Eqs. (36) and (37) define their angular dependencies for bi-layers and axons, respectively. Experimental data (see below) show that the anisotropic contributions to MR signal comes mostly from THBs with lifetimes *τ* _*A*_ in the range of tens nanoseconds. In this interval, the general Eqs. (6) and (7) can be approximated by their asymptotic forms:

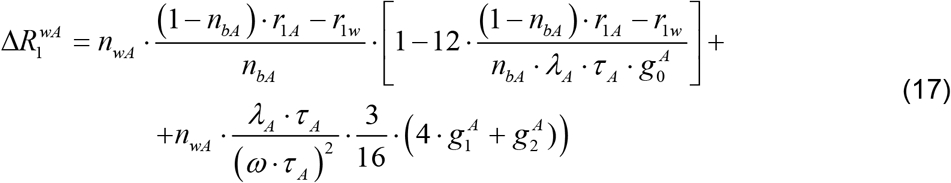

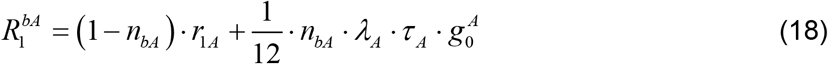

with angle-dependent parameters 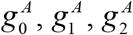 defined by Eqs. (36) and (37) for bi-layers and axons, correspondingly. Since for typical cases (*nbA* · *kA*) ≫[(1− *nbA*)· *r*1*Ar*1*w* ] (see experimental results below), the contributions of the second and the third terms in Eq. (17) are small. Thus, the anisotropy of water signal 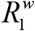 is mostly defined by the anisotropy of *r*_1*A*_, i.e., by the bound spin-spin interactions in the bi-layers. Herein, in analyzing experimental data, we will use results for *r*_1 *A*_ obtained in the framework of the LDM (Eq. 19 for bi-layers and Eq. 22 for axons in [42]) as reproduced herein in the Appendix 3, Eqs. (38) and (39).

### Basic THB Model of the Transverse (T2) Relaxation in CNS

It is usually assumed that the *b*-protons have very short, on the order of tens microseconds T2 relaxation times [51-53], *r*_2 *j*_ ≈ 10^4^ − 10^5^ sec^−1^. Hence, their contribution to the signal in the millisecond range (where most MRI experiments are conducted) is very small and will not be considered herein. These terms, most likely, are the sources of the signals detected by the so-called ultra-short TE techniques (see, e.g., [52, 54-57]). However, it is important to note, that the amplitude of the water signal, 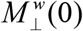, depends on flip angle *θ* in a regular manner:

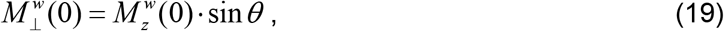

while the amplitudes of *b*-signals, 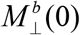, depend on *θ* in a more complicated manner modulated by the value of *r*_2 *j*_ (see Eqs. 9,10, and 19 in [58]).

In the framework of the Basic THB Model with two *b*-pools (as discussed above for the T1 relaxation), the result for the transverse component of the water signal, in the main approximation with respect to the small parameters 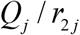, can be presented as follows:

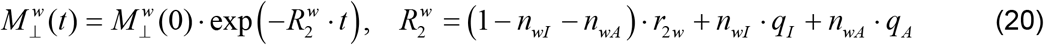

where

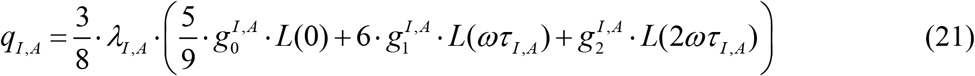

with the coefficients *g* ^*I, A*^ defined in the Appendix 1 by Eqs. (35)-(37).

The dependence of the isotropic part of 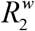 in Eq. (20), 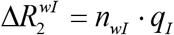, on the THB network lifetime is illustrated in Figure 3 reveals that the isotropic part of 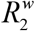 becomes *B*_0_ -independent and for *ωτ* _*I*_ > 6 approaches its asymptotic behavior:

**Figure 3.**
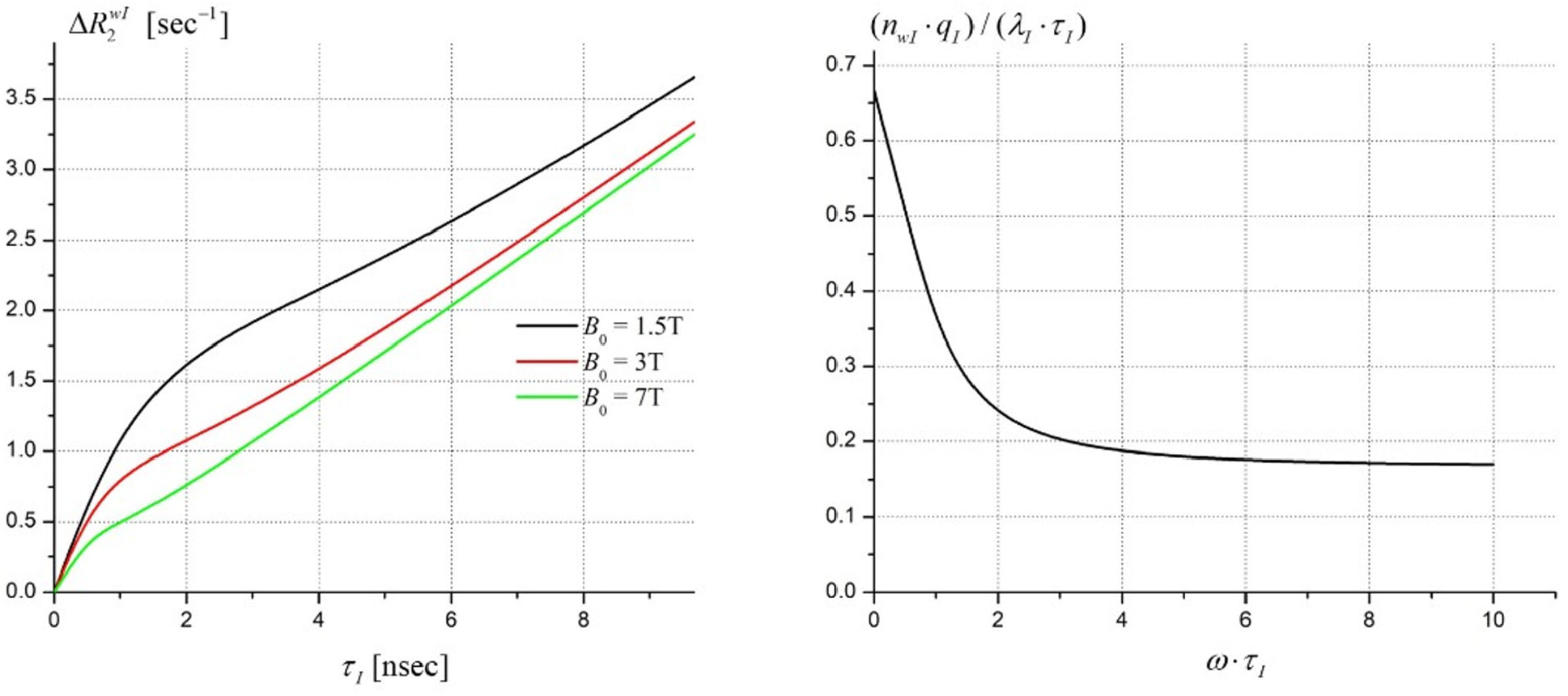
General properties of the isotropic part 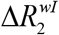. Left panel: 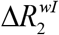 as a function of the lifetime *τ* _*I*_ for *λ* = 20 (sec· nsec)^−1^, *n*_*wI*_ = 0.1, and different external magnetic fields *B*_0_. Right panel: the characteristic behavior of 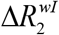 as a function of *ωτ*_*I*_.

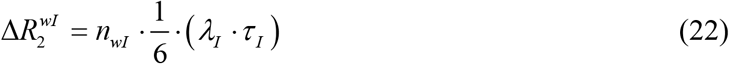

The angular dependence of the anisotropic part of 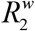 in Eq. (20), 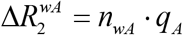, for the axonal structure is illustrated in Figure 4.

**Figure 4.**
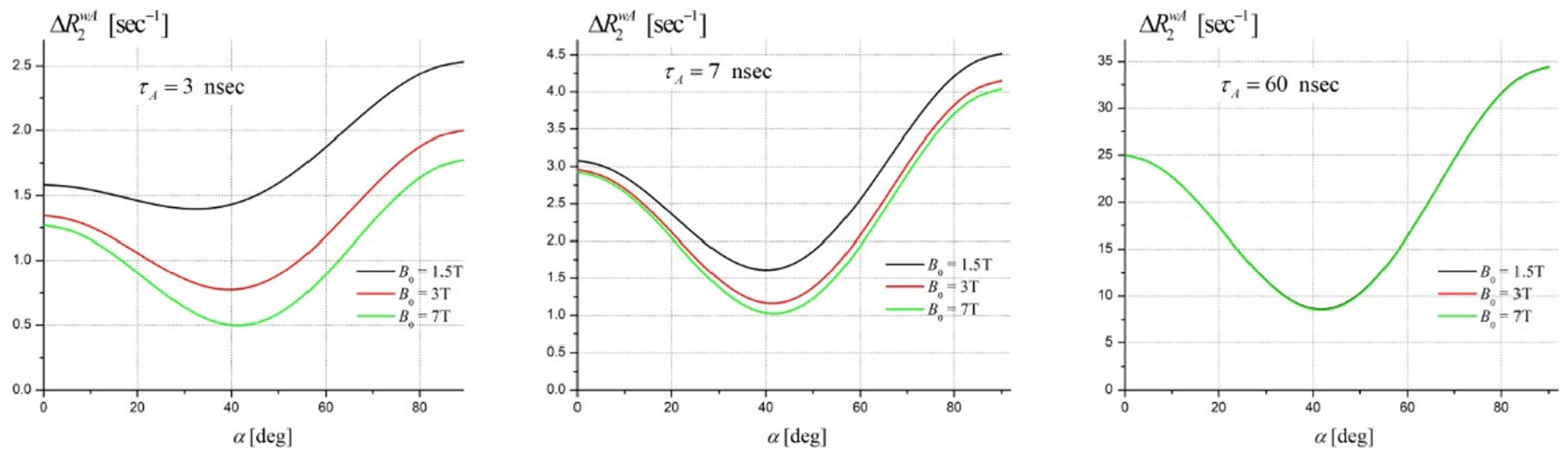
The anisotropic term 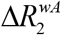 in Eq. (20) in the axonal structure. The plots illustrate the angular dependence of 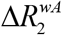 for *λ* = 20 (sec· nsec)^−1^, *n*_*wA*_ = 0.1, different correlation times *τ* and magnetic fields *B*_0_.

For long correlation times *τ*_*A*_ (*ωτ*_*A*_ >>1), the anisotropic term 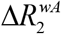 becomes independent of magnetic field *B*_0_ (right panel in Figure 4), and for axonal structures can be presented by the following asymptotic expression:

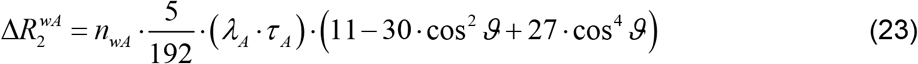

### Basic THB Model Validation and Parameters’ Estimation in CNS

Data were obtained from previously published paper by Wallstein, et al [43]. Per their description: “Fresh pieces of porcine spinal column from a local slaughterhouse (postmortem interval 2–3h) were fixated in 4% PFA/PBS solution. A piece of spinal cord from segment L5/6 (lumbar vertebrae) was carefully dissected and a part of the WM region was placed in an NMR tube embedded in Fomblin. Finally, the tube was placed in the central region of a Helmholtz coil. The setup allows remote reorientation of the coil and sample around the cylindrical axis of the coil, which runs through the isocenter of the magnet. The sample orientation was varied between angles of 0° (corresponding to a parallel alignment with the external field B0) and 90° (corresponding to a perpendicular alignment with the external field B0). Measurements were performed with a total of 11 rotation angles. All experiments were performed on a MAGNETOM Skyra^fit^ 3T scanner (Siemens Healthineers, Erlangen, Germany).” The authors provided results for several T1 measurements of which we use Inversion Recovery (IR) that were performed with soft, adiabatic inversion pulses (BIR-4, *τ* _*p*_ = 5 msec). Datasets consisted of 23 logarithmically spaced inversion times (770 μsec ≤ TI ≤ 10 sec, *TR* = 13 sec). Further experimental details can be found in [43].

For our data analysis we used IR data acquired from two samples (A and B) at two temperatures (22°C and 36°C). Since *b*-protons have very short, on the order of tens microseconds, T2 relaxation times [51], the soft inversion pulse excited only water protons and had very little effect on the initial *b*-protons magnetization (i.e.,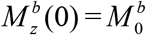). Hence, neither Eq. (5) nor the contribution described by the last term in Eq. (4) were used in the data analysis.

An important feature of the THB theory is the prediction of anisotropic behavior of T1 and T2 relaxation. While the THB theory predicts MR signal dependence due to the interactions between water and bound protons (*w-b*), the results also depend on the *b-b* relaxations defined by the relaxation parameters *r*_1*I*_, *r*_1*A*_. Herein, in analyzing experimental data, we will use results for *r*_1*A*_ obtained in the framework of the LDM (Eq. 19 for bi-layers and Eq. 22 for axons in [42]) as reproduced herein in the Appendix 3, Eqs. (38) and (39). The analysis of Eq. (39) for axons reveals that at 3T for parameter Ω > 10 (Ω= *ω* ·*τ* _*d*_, *τ* _*d*_ is the characteristic lateral diffusion time), the angular dependence of *r*_1 *A*_ only slightly changes with Ω and can be parametrized by the following phenomenological function

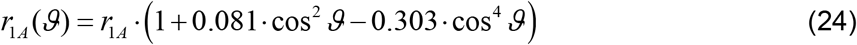

with the angle-independent parameter *r*_1*A*_.

In present study, we use the Bayesian probability approach developed by Larry Bretthorst [59, 60]. For each set of experiments with different rotation angles, data was fit by using the joint probability analysis, in which all the THB model’s parameters (*λ*_*I, A*_, *n*_*wI, wA*_, *n*_*bI, bA*_, *τ* _*I, A*_, *r*_1*I, A*_, *r*_1*w*_) were assumed the same (as they reflect biological tissue MR properties) and only amplitudes of the signals, 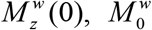, corresponding to different rotation angles were allowed to vary independently. For the relaxation of pure water, *r*_1 *w*_ = 0.24 sec^−1^, is assumed [35]. Using Eq. (31), and the fitting result for the parameter *λ*, we can also calculate an average dipole-dipole distance *d* [Å] in THBs:

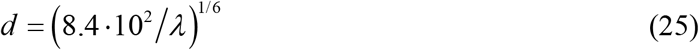

where *λ* in (sec· nsec)^−1^.

Typical result of the Basic THB model fitting to experimental data is shown in Figure 5 that demonstrates a rather good fit with residual not exceeding 1% at any data point. The post-fitting calculations based on Eqs. (14) reveal a rather significant (∼ 10%) *w-b* components’ contribution to the signal in the interval of the inversion times up to 3 seconds. Below we use notations Iso and Aso for the Isotropic and Anisotropic pools.

**Figure 5.**
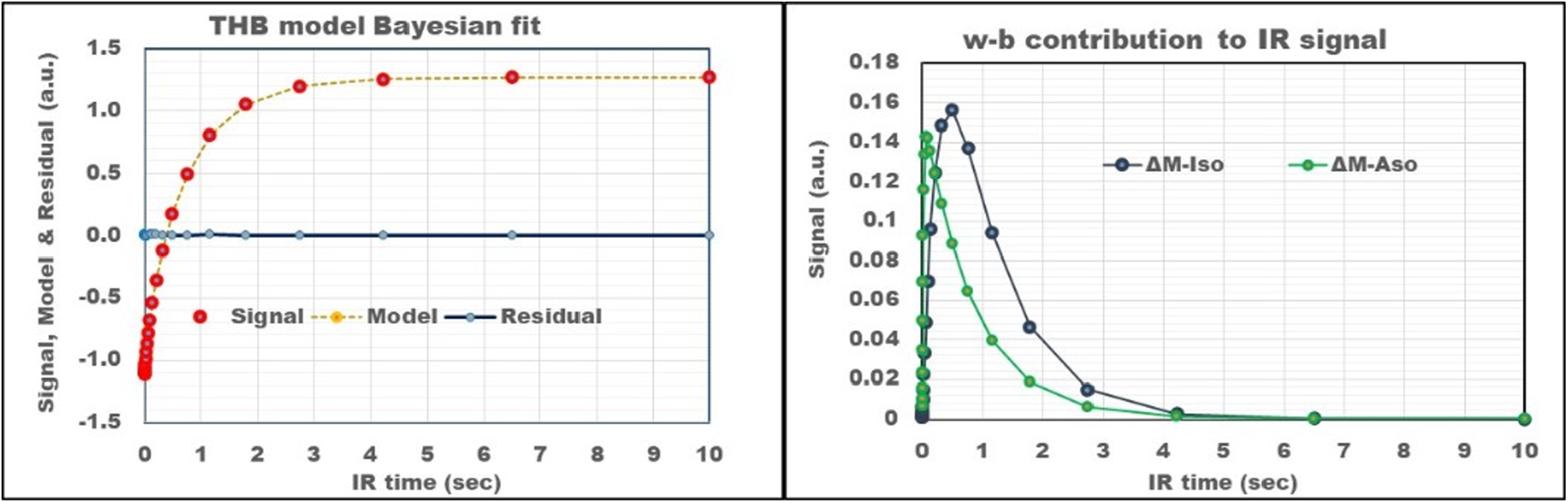
THB model fitting to Inversion Recovery data. <u>Left panel</u>: Example of inversion recovery data obtained from sample A at physiological temperature 36°C, and oriented at 90° to **B**_0_. Red circles – actual signal; orange circles and line – fit the THB model in Eq. (4) with 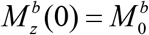(no *b*-protons excited by RF pulse) to experimental data; blue connected circles – residual of fitting. <u>Right panel</u>: Contribution of isotropic and anisotropic *w*-*b* terms in Eqs. (14) to the IR signal model.

A summary of the basic biophysical THB model parameters obtained from all datasets is presented in Figure 6 and Figure 7. In general, the results show consistency between two samples A and B, but also slight differences between measurements at two temperatures (36°C and 22°C). This is expected since temperature affects the dynamic interactions between water and bound protons. An important finding is five-to-ten-fold differences between concentrations of isotropic and anisotropic transient hydrogen bonds.

**Figure 6.**
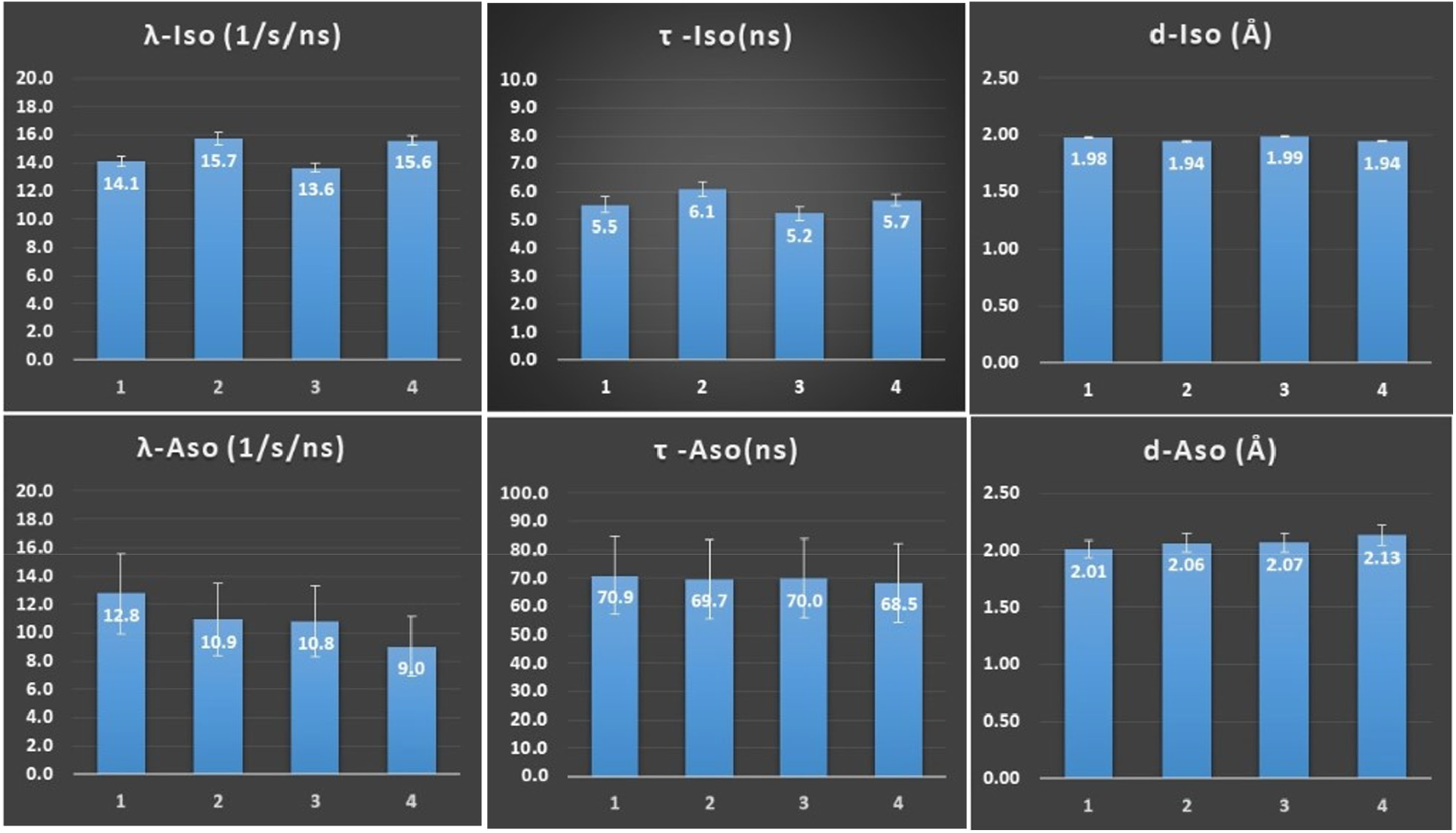
Parameters of the quantum dipole-dipole interactions in the THBs obtained from inversion recovery experiments. The parameter *λ* characterizes the strength of interactions, parameters *τ* – their lifetimes, and parameters *d* – the distances between *w*- and *b* protons in THBs. Bars show mean values of the Bayesian probability distributions of the estimated THB model parameters; error bars correspond to the STD of Bayesian probability distributions (representing errors in parameters’ estimates). Notations: (1) Sample A, 36°C; (2) Sample A, 22°C; (3) Sample B, 36°C; (4) Sample B, 22°C.

**Figure 7.**
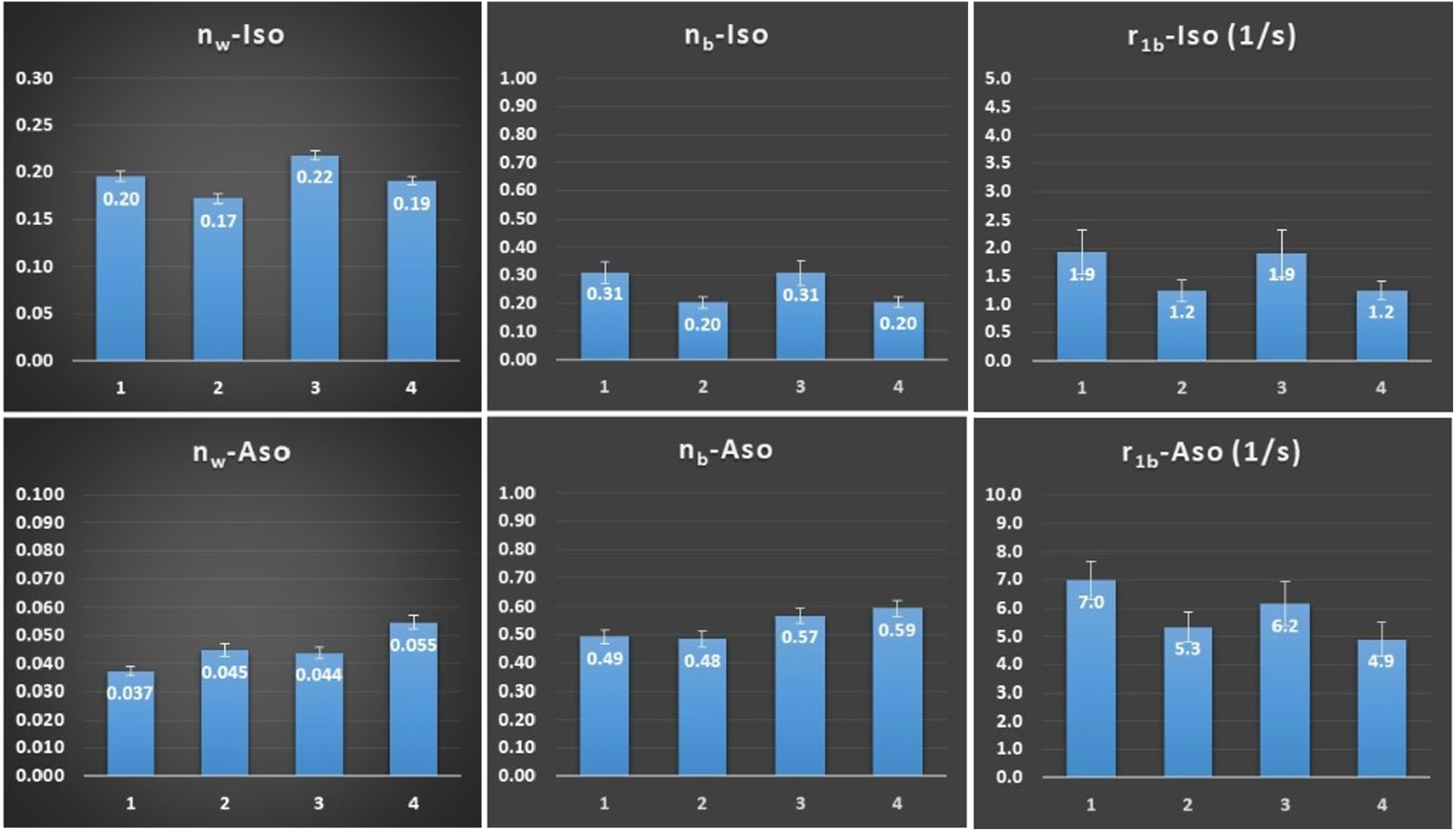
Parameters characterizing relative concentrations of water protons (*n*_*w*_) and bound protons (*n*_*b*_) participating in the THBs, as well as the bound protons’ relaxation rate parameters (*r*_1 *b*_) obtained from the analysis of the IR experiments. Bars show mean values of Bayesian probability distributions of estimated THB model parameters; error bars correspond to STD of Bayesian probability distributions (representing errors in parameters’ estimates). Notations: (1) Sample A, 36°C; (2) Sample A, 22°C; (3) Sample B, 36°C; (4) Sample B, 22°C.

By making use of parameters in Figure 6, Figure 7, and Eqs. (6), (7), (11), we can evaluate values of water 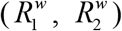 and bound protons, 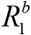, relaxation rate parameters and their angular dependences. The result for the sample A at 36°C is presented in Figure 8, showing only slight anisotropy of the water 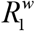 relaxation rate parameter, but significantly higher values and anisotropy of the water 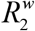 and the bound protons’ 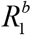.

**Figure 8.**
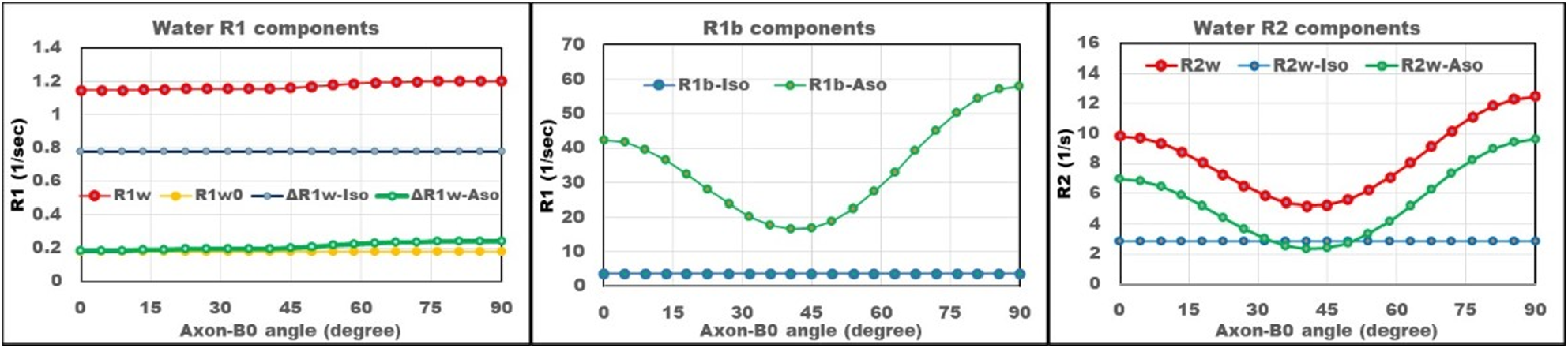
Example of water 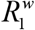 (left panel), bound protons 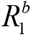 (middle panel), water 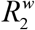 (right panel) relaxation rate parameters as functions of the angle between sample orientation and external magnetic field **B**_0_ for the sample A at physiological temperature 36°C. The curves are calculated based on Eqs. (6), (7), (11), (20) and the biophysical THB model parameters presented in Figure 7. Red symbols – total relaxation rates, green symbols – contributions of anisotropic THB pool, blue symbols – contributions of isotropic THB pool, and yellow is contribution of *r*_1 *w*_.

## DISCUSSION

Inherent differences in MR signal relaxation properties between diverse biological tissues and their conditions (i.e., healthy vs. diseased) result in multiple applications of MRI in biology and medicine. Since inception of MRI, numerous papers have been devoted to understanding basic mechanisms relating MR signal relaxation properties to underlying microstructure of biological tissues. Such a knowledge allows improved MRI techniques better outlying tissue anatomy (e.g., gray matter vs. white matter) and pathology (e.g., tumors, lesions, etc.). Hence, theoretical models describing relaxation properties of MRI signal should account for salient features of tissue microstructure related to specific questions that MRI experiments are trying to address.

In this paper we explore and further develop the Transient Hydrogen Bond (THB) model [35] that accounts for the salient features of the quantum dipole-dipole interactions between protons of the water molecules (so called “free” protons) and protons belonging to the tissue-building molecules (so called “bound” protons). The former, “free” protons of water molecules, randomly walk (diffuse) through multiple tissue environments, spending a portion of travelling time being transiently attached by hydrogen bonds to macromolecules in lipid heads of cellular and myelin bilayers, where they experience quantum spin exchanges with “bound” protons, thus causing relaxation of the MR signal.

Our results show that the MR signal generated from such a dynamically interacting system requires description in terms of several subsystems of THBs (at least, two in the introduced Basic THB model) resulting in three exponential components with their relaxation properties defined by quantum interactions as described by Eqs. (4) and (5) for T1 signal and T2/T2* signals. Thus, analyzing experimental measurements of MR signal relaxation in the framework of the THB theory can provide a specific information on the organization of cellular membranes and myelin in biological tissues.

Bound protons interacting with water protons represent a *sub-population of protons in the hydrophilic heads and proteins*. It is important to note that not all protons in macromolecules, but only those *structurally available* for close transient contact with free water, contribute to the magnetic spin exchange, hence, to the relaxation properties of measured water MR signal. The magnetic properties of these bound protons are significantly altered by interactions with the dynamic and constantly renewing pool of water molecules within the available space at the periphery of the bilayer structure. Conversely, water protons are less affected, as their interaction with bilayer hydrophilic heads is transient, representing only a portion of their diffusive pathway.

In this paper, by analyzing previously published IR data obtained by Wallstein, et al [43] from porcine spinal column, we have established the main parameters of the Basic THB model. Our data has proven the presence of two types of transient hydrogen bonds with the properties presented in Figure 6 and Figure 7. The first, Iso-type, is characterized by the isotropic distribution of dipole-dipole connections with THB lifetime (*τ* _*I*_) about 5-6 ns and strength of connections described in the THB model by the parameter *λ*_*I*_ of about 14-15 (sec· nsec)^−1^. The second, Aso-type, is responsible for the anisotropic properties of MR signal and is characterized by the THBs forming orientation order with direction of w-b connections perpendicular to the bi-layers surfaces. This group of THBs have significantly longer lifetime (*τ* _*A*_) about 70 ns and the slightly smaller strength of connections described by the parameter *λ*_*A*_ of about 9-12 (sec· nsec)^−1^. For both the THB groups, the distances between dipoles in *w*-*b* connections are quite similar - about 2 Å. It is important to note that the presence of the THBs with long lifetimes identified by application of the THB model to experimental data in porcine spinal cord is consistent with the results of Koenig [61], who reported that the immobilized protein solutions can trap solvent water molecules for times from 40 nsec to up to 1 μsec. This potentially suggests that the long-time THBs can be associated not only with macromolecules in the lipid heads of cellular and myelin bilayers, but also with the structurally organized proteins forming cytoskeleton (such as microtubules occupying axons and dendrites).

The dynamic pathways of water and bound protons are quite different. The water protons diffuse throughout the cellular bed, while the bound protons, attached to long macromolecules forming cellular membranes (bi-layers), participate in the lateral diffusion of this macromolecules as described by the LDM [42]. The dipole water-water interactions outside the THBs give very small contribution to T1 relaxation (*r*_1_*w* ∼ 0.24 sec^−1^), while the dipole-dipole *b*-*b* interactions, due to the lateral diffusion, lead to substantial contributions to their T1 relaxation (*r*_1 *b*_ -Aso is about 5 − 7 sec^−1^, and *r*_1*b*_ -Iso is about 1 − 2 sec^−1^), as demonstrated in Figure 6.

The important parameters characterizing contribution of THBs to T1 IR signal are the parameters *n*_*wI*_ and *n*_*wA*_ representing fractions of water protons of type Aso and Iso participating in the Aso and Iso transient hydrogen bonds at any given moment. These fractions are very different – they are about 17-22% for Iso THBs and only 4-5% for Aso THBs. The fractions of bound protons, *n*_*bI*_ and *n*_*bA*_ in the *w*-*b* connections are much bigger: about 50-60% for Aso THBs and about 20-30% for Iso THB. We can hypothesize that the bound protons contributing to the isotropic subtype of THBs are located mostly at the periphery of the hydrophilic heads of the bi-layers, directly adjacent to the freely diffusing water. Hence, their structure is mobile and only loosely oriented. On the contrary, the bound protons contributing to the anisotropic subtype of THBs, reside deeper within the hydrophilic heads with their orientation exhibiting greater structural rigidity (the THB orientation order), resulting in fewer potential interactions with water protons.

The idea of the *THB order* introduced in our theoretical approach is consistent with the results of Ohto et al [62] on the structure and orientation of water at zwitterionic phosphatidylcholine. The authors demonstrated that the lipid carbonyl groups served as efficient hydrogen bond acceptors allowing the water hydrogen bond network to reach, with its (*up*-oriented) O–H groups, into the headgroup of the lipid.

Our analysis shows that water interactions with both, Iso and Aso THBs, contribute significantly to the IR signal, as illustrated in Figure 5, especially at IR times below 2-3 seconds, where these contributions account for more than 10% of the total IR signal. Importantly, these contributions are not monotonic (second and third terms in Eq. (4)), starting from zero right after inversion pulse, reaching peak at IR time

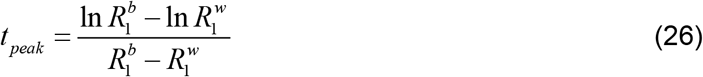

and then slowly decaying as 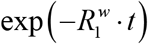. Positions of these peaks are closer to the origin for the Aso THB as its relaxation rate parameter (ranges between 15 and 60 sec^-1^) is much bigger than for the Iso THB with the relaxation rate parameter of about 4 sec^-1^ (Figure 8, middle panel). The relatively smaller contribution of long spin-exchange component to IR signal explains why the relative uncertainties (STD in Figure 6 and Figure 7) for the parameters defining component with long spin-exchange times are much bigger than for the parameters defining short spin-exchange component.

While both, Aso- and Iso-THB networks contribute significantly to the IR signal, their contributions to the relaxation rate parameters are quite different, as illustrated in Figure 8. The Iso-THB network gives major contribution to the T1 signal relaxation (Figure 8, left panel) but small to T2 relaxation (Figure 8, right panel). On the contrary, the Aso-THB network gives very small contribution to the T1 relaxation but very significant to the T2 relaxation. Such differences in Iso- and Aso-THBs contributions to the relaxation parameters are consequences of the differences in their lifetimes. While THBs with lifetimes *τ* _*I*_ around 3-6 nsec, are favorable for T1 relaxation (Figure 1), their contributions to T2 relaxation is relatively small in this lifetime range (Figure 3). Conversely, due to their long lifetimes *τ* _*A*_, around 70 nsec, Aso-THBs give dominant contribution to T2 relaxation, as described by Eq. (23), but very small to T1 relaxation, Eq. (17).

The above-discussed differences in Aso- and Iso-THB lifetimes explain why the anisotropy of water R1 relaxation rate parameter is very small (green symbols in the left panel of Figure 8) but quite significant for the water R2 relaxation rate parameter (green symbols in the right panel of Figure 8). It is also important to note that due to the long lifetime of Aso-THBs, the anisotropy of 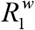 is mainly determined not by THBs, but by the anisotropy of the relaxation term *r*_1 *A*_ defined by the *b*-*b* interactions in the bilayers (Eq. (17)).

### THB model explains cellular specific contribution to the transverse (T2/T2*) relaxation of MR signal

The THB model suggests that the contribution of hydrogen bonds to the transverse relaxation, Eqs. (20)-(23), is proportional to the fractions of water protons in the transient hydrogen bond state in the lipid heads of bilayers. Hence, the parameters *n*_*wI*_ and *n*_*wA*_ are proportional to the density of bilayers. In WM, where bilayers are wrapped around axons, forming a myelin sheath, these parameters are proportional to the myelin density. This is consistent with numerous reports that established strong associations between R2* relaxation in WM and tissue myelin content [29, 63-67].

While the longitudinal relaxation rate decreases with magnetic field increases, the transverse relaxation rate increases at large magnetic fields. From this perspective, the THB model explains only a part of the transverse relaxation as the THB contribution remains practically field-independent and even slightly decreases with *B*_0_ increases (see Figure 4). The remaining part is most likely related to the presence of iron and magnetic susceptibility effects (see recent publications [65-68] and references therein).

The strength of the THB interactions defines their contribution to the T2 relaxation of water signal. For an example, in Figure 4, their contribution to 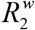 is in the range of 20 sec^-1^. For parameters corresponding to the porcine spinal cord, the THBs contribution to 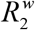 is about 10 sec^−1^. These values of 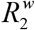 are consistent with the values usually measured in WM. Indeed, Bartzokis [45] reported 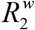 of 14.5 sec^-1^ at 0.5T and 15.6 sec^-1^ at 1.5T, Whittal et al [69] reported 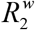 of 12.9 sec^-1^ at 1.5T, Gelman et al [70] and Wang et al [68] reported 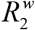 of 18 sec^-1^ at 3T. Similar results were also reported by van Gelderen, et al [19] for brain regions with low iron content. These in vivo data are consistent with ex-vivo results of Stanisz et al [71] for fixed tissue: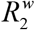 of 13.9 sec^-1^ at 1.5T and 14.5 sec^-1^ at 3T. While it is known that the tissue R2 can be affected by the presence of iron [45, 70], especially at high fields [19, 68], the low field data suggest the presence of the iron-independent component. Also, a value of 14 sec^-1^ was obtained at 3T for R2t* (a tissue specific subcomponent of R2*) after separating contributions to R2* from heme and non-heme iron and macroscopic magnetic field inhomogeneities [72].

### Transient Hydrogen Bond Order is a primary source of the longitudinal (T1) and transverse (T2/T2*) MR signal relaxation anisotropy in WM

We first note that in the framework of the THB model the formation of MR signal is described by the transient dipole-dipole interactions happening at the nanosecond time scale. From this perspective, THB-induced changes in the MR signal cannot be refocused by the RF pulses of microsecond-to-millisecond duration used in traditional MR experiments. Hence, the THB mechanism will provide identical contribution to both, T2 (i.e., spin echo) and T2* (i.e., free induction decay or gradient recalled echo) signals. Accordingly, the discussion below can be equally attributed to both T2 and T2* considerations.

Following an initial demonstration of the transverse R2* signal anisotropy in the neuronal tissue, obtained by the actual brain rotation by Wiggins et al [20], field-to-fiber measurements by Cherubini et al [21], and the prediction of the axonal structure-specific anisotropic behavior of the transverse MR signal phase in WM [73], numerous papers studied different aspects of these anisotropic phenomena in WM. While several *phenomenological* models were proposed to describe R2* [22, 26, 27, 29, 74-78] and R1 [33, 79] relaxation anisotropy, the THB model offers the specific biophysical mechanism explaining practically all existing to-date experimental data on MR signal relaxation anisotropy in the mammalian brain.

Indeed, the introduced herein Basic THB Model explains major features of the MR signal anisotropy in the brain observed in the T2* relaxation: higher R2* for axons perpendicular to the external magnetic field **B**_0_ as compared with axons parallel to **B**_0_ and a broad minimum around 40-50°. The difference in R2* between perpendicular and parallel axonal orientations (denoted below as ΔR2*) was found about 3 to 5 sec^-1^ in [22, 74, 75]. Importantly, these features did not depend on the strength of magnetic field **B**_0_ that is consistent with the results reported by Oh et al

[27]. As demonstrated in [27] (7T whole body MRI, formalin fixed human brains), extracting tissue iron did not reduce the orientation-dependent R2* contrast in corpus callosum (ΔR2*=51.3-47.5=4.0 sec^-1^ before extraction, and ΔR2*=43.6-39.2=4.5 sec^-1^ after extraction at 21°C). This fact suggests that iron is not the origin of the anisotropy of R2*, though affecting its values. Their result also did not depend on the fixation status which was consistent with increased postmortem R2* in WM fibers for both parallel and perpendicular measurements as compared with in vivo data (but with ΔR2* remaining at about 4 sec^−1^) reported by Lenz et al [80].

Sati et al [26] provided in vivo data from marmoset brains at 7T showing that R2* relaxation in WM was overly sensitive to the fiber orientation relative to **B**_0_. Depending on a WM fiber track, the ΔR2* ranged from 5 sec^-1^ to 14.7 sec^-1^ in the optic radiation. These results are in agreement with the results obtained by Wiggins [23], who reported the difference ΔR2*=14.8 sec^-1^ for some fibers (7T, the human brain). Importantly, the data in [26] and [23] have been obtained in vivo using the actual brain rotation.

Detail measurements of R2* anisotropy in fixed rat brain at 9.4T MRI (actual brain rotation) was reported by Rudko et al [29]. They demonstrated the angular variation of R2* with difference of 4 sec^-1^ between perpendicular and parallel directions and a broad minimum around 40°.

It is also interesting to mention data obtained by Weber et al [75] for neonate brain, showing a non-monotonic angle dependence but with the practically identical (8.6 s^-1^) values of R2* for the parallel and perpendicular orientations and with the minimum (8.3 s^-1^) at about 40° from the parallel orientation. Though we should note that these data have been obtained by measuring the fiber-to-field angle rather than the direct rotation, hence they can be affected by a potentially different structure of white matter bundles. Also, smaller values of R2* are consistent with prior R2* measurements in neonatal brain [81].

Similar to T2/T2* signals, the anisotropy was also reported in the T1 signal by Kauppinen et al [33, 79]. Recent detail studies by Wallstein, et al [43] demonstrated that the specifics of this anisotropy quite significantly depend on the method used for T1 measurements. Specifically for IR experiment, they reported changes in the T1 angular behavior as a function of the initial inversion time used for data analysis and attributed this effect to a multi-component contribution to the IR signal. Though, no specific mechanism was proposed to explain this effect. Herein we demonstrated that the IR signal time behavior and its anisotropy (exemplified by data in the porcine spinal cord) can be very well described in the framework of the Basic THB theory accounting for contribution of two types of THBs – one with isotropic distribution of *w*-*b* connections (Iso-THBs), and another with anisotropic distribution of *w*-*b* connections (Aso-THBs).

Data in Figure 9 demonstrate that ignoring contributions from THBs to IR signal and, instead, using a simplified model with a single exponential, would lead to significant acquisition-dependent errors in quantifying water T1 signal. An example shown in the left panel of Figure 9 demonstrates the dependence of the R1 relaxation rate measurements in the porcine spinal cord by means of the IR experiment on the signal sampling interval. One can see that when initial inversion time changes from 1 msec to 1 sec (keeping the final sampling time at 10 sec), the R1 decreases by 30%. The best match with correct water signal 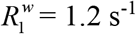 for perpendicular orientation and 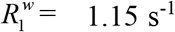 for parallel orientation is achieved when IR signal is sampled between 1 and 10 seconds. Data in the right panel of Figure 9 explain the variation in the R1 signal anisotropy in the framework of the mixed THB model. The important issue here is a very small and almost monotonic variation in 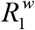 with the angle between **B**_0_ and axonal direction, as shown in Figure 8. At the same time, there is a very strong angular dependence of the relaxation rate parameter 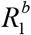 -Aso (Figure 8). Hence, if data are fitted using a single exponential model, the fitting parameter R1 (in this case, it should be called an *apparent* R1, 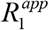) actually represents a mixture of 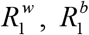 -Aso and 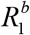 -Iso. The data in the right panel of Figure 9 show examples of 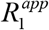 calculated for the mixed signal model according to the equation:

**Figure 9.**
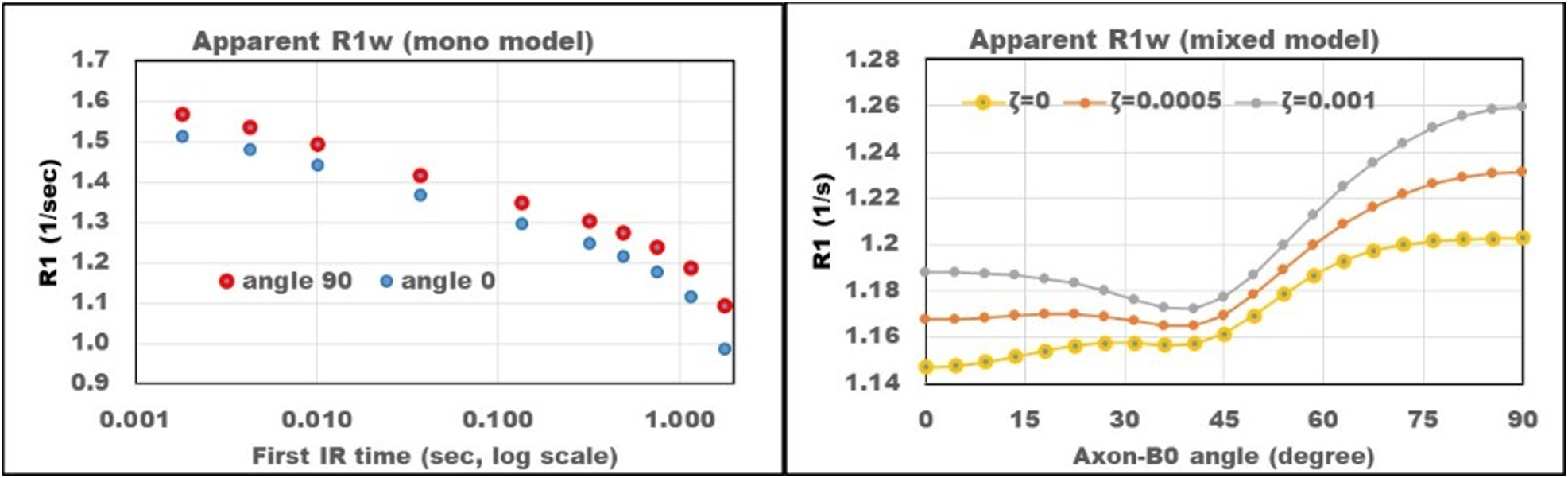
Example of IR data analysis using a model with a single exponential component. Data represents inversion recovery signal obtained from sample A (Wallstein, et al [43]) at physiological temperature 36°C. Left panel demonstrates the dependence of the R1 relaxation rate on the initial IR sampling time for two samples oriented at 0° and 90° to **B**_0_. Data in the right panel explain thevariation in the R1 signal anisotropy determined by means of the mixed model, per Eq. (27). The result shows that even for a very small mixing parameter ζ, on the order of 0.1%, the character of the angular dependence changes from practically monotonic increase for ζ=0, to a behavior with a broad minimum around 40°.

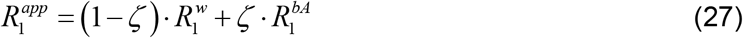

where 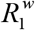 is defined by Eq. (6), and 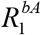 (i.e., 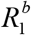-Aso) is defined by Eq. (18). In the Eq. (27), the contribution from 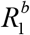-Iso was omitted because it is much smaller than the 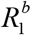-Aso and would give a minimal contribution at these very small mixing rates. The result shows that even for a very small mixing parameter *ζ*, on the order of 0.1%, the character of the angular dependence changes from practically monotonic increase for *ζ* = 0 to a behavior with a broad minimum around 40°. Such a behavior is consistent with the results of T1 measurements by Kauppinen et al [33, 79] reporting a broad peak centered around 40-50° at 3T and 7T, and a smaller R1 for WM fibers parallel to the magnetic field **B**_0_ as compared to the fibers perpendicular to **B**_0_. Since the authors of [33, 79] used the IR-type method for data acquisition with a single exponential model for data analysis and a rather restricted IR measurement times, their data can be explained by a mixed model for the defined by Eq. (27) and presented in Figure 9.

Recent multi-center multi-vendor study [82] reported on the inherent uncertainty in the T1 measures of water signal across independent research groups. These uncertainties are not surprising due to the enormous complexity of biological tissues, where hundreds (if not thousands) cells and subcellular structures are usually present in each imaging voxel. This microstructural complexity varies across different biological structures and even within similar structures that change with biological tissue age and diseases-related pathology. As demonstrated in our study, the cause of T1 relaxation can be well explained by the interaction between water molecules and cell-building materials, leading to multiexponential behavior of T1 signal. On the contrary, by analyzing data in the framework of a single exponential model (as was the case in [82]) would always provide estimates of T1 relaxation time parameter that depend on the parameters of the pulse sequence used in the experiment. Even though the Inversion Recovery (IR) is considered a gold standard for measuring T1 relaxation, example in Figure 9 demonstrates a variation in the T1 measurements of about 30% to 40%, depending on the IR sequence parameters and sample orientation. A robust estimation of relaxation parameters requires biophysical modeling that captures salient features of MR signal as demonstrated in this paper by means of the Basic THB model.

### In Summary

In this paper we have introduced and validated Basic Transient Hydrogen Bond model that provides quantitative evaluation of MR signal relaxation in CNS. The model explains MR signal relaxation as a result of magnetization exchanges enabled by quantum dipole-dipole interactions withing Transient Hydrogen Bond matrix comprised of water and lipid protons residing within hydrophilic heads of lipid bilayers forming cellular and myelin membranes. We demonstrated the presence of two types of THBs with different lifetimes. One – with the lifetime of a few nanoseconds – is mostly responsible for the longitudinal relaxation of MR signal. The second – with the lifetime in the range of tens nanoseconds – mostly contributes to the transverse relaxation and is responsible for the anisotropy of both, longitudinal and transverse relaxations. The ability of the Basic THB model to differentiate distinct THB Matrix components based on their MR signal relaxation properties can be fundamental for identifying specific pathological changes, thus enhancing disease specificity on MRI scans.

## Appendix 1: General definition of the THB model parameters

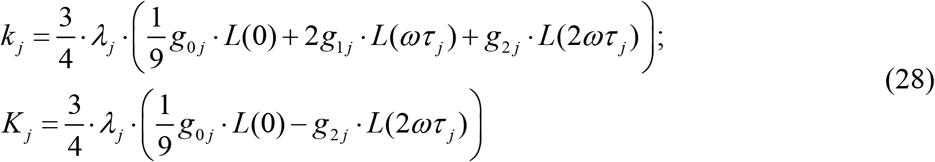

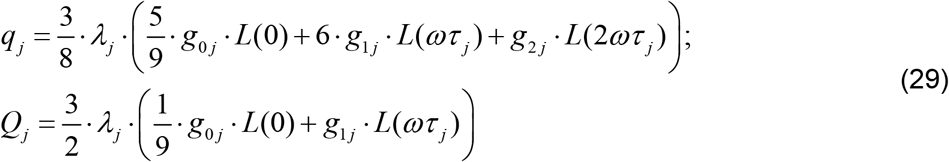

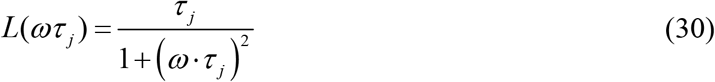

*ω* = *γ B*_0_, *τ j* is the *j*^th^ THB lifetime. Recall that per THB theory [35], the correlation functions are described by the stretched spectral density function accounting for the presence of multiple THB lifetimes. Since our new approach already accounts for multiple THB lifetimes, in what follows we will use the standard Lorentzian form in Eq. (30) for each individual THB type.

The parameters *λ j* characterize the strength of quantum dipole-dipole spin exchange interactions between water protons transiently trapped in lipid heads and their lipid-bound counterparts to the MR signal relaxation:

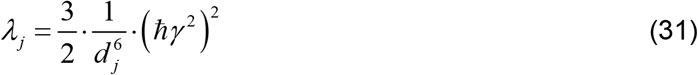

where *d j* is a distance between *w-b* protons in the THB.

The angular dependences of the diagonal and cross-relaxation parameters in Eqs. (28) are defined by functions

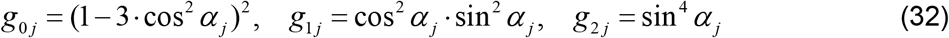

where *α j* are the angles between corresponding dipole-dipole *w-b* connection and direction of the external magnetic field *B*_0_.

## Appendix 2: Definition of model parameters of the Basic THB T1 and T2 models

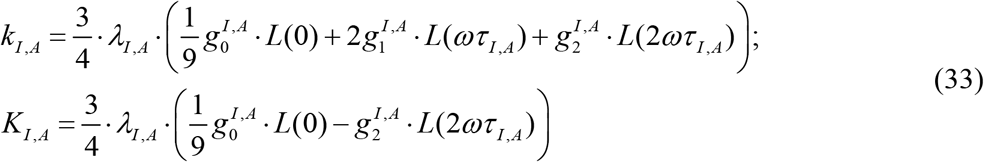

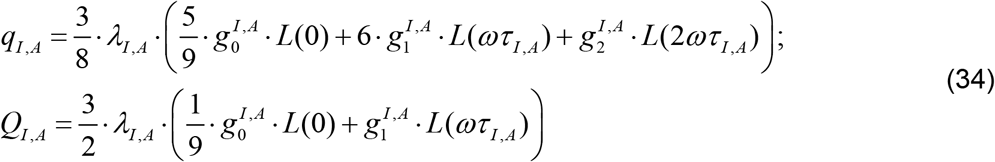

The structure of coefficients *g* for different distributions of *w-b* orientations in THB was discussed in details in [35]. In particular, the coefficients *g* ^*I*^ of isotropic network, obtained by averaging Eqs. (32) with the isotropic distribution function, are:

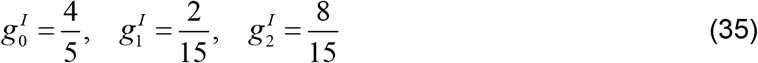

For individual bilayers, the anisotropic coefficients *g*^*A*^ are

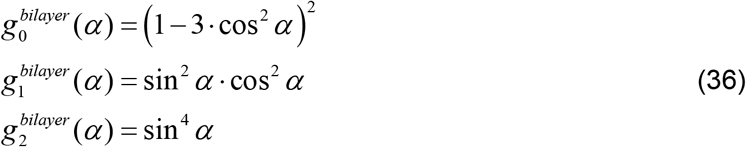

where *α* is the angle between bilayer normal and the external magnetic field For the white matter, and specifically, for bilayers covering individual axons (myelin sheath), coefficients *g* ^*A*^ are obtained by averaging Eqs. (36) with the distribution function corresponding to the orientation order of the dipole-dipole *w-b* connections oriented perpendicular to the surfaces of bi-layers surrounding axons. The result is:

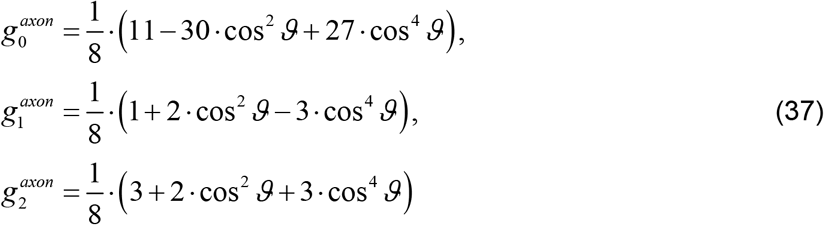

where *ϑ* is the angle between direction of the axonal bundle and

## Appendix 3: Longitudinal relaxation parameters due to the bound proton *b-b* interactions

Herein we present the results previously obtained in the framework of the LDM (Lateral Diffusion Model) [42]. The following equations define *r*_1 *A*_ for bi-layers and axons:

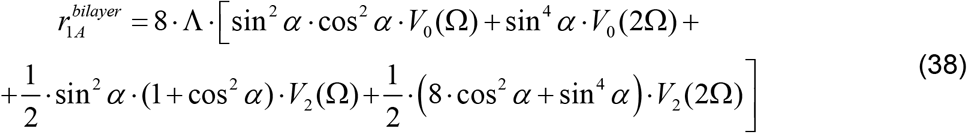

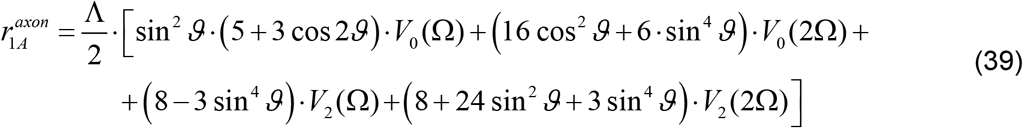

where Ω= *ω* ·*τ* _*d*_, *τ* _*d*_ is the characteristic lateral diffusion time; the functions *V*0,2 (Ω) and parameter Λ are defined in [42]. Note that for Ω> 50 the following asymptotic expressions provide accurate values of the functions *V*0,2 (Ω) : *V*0 (Ω) ≈ 0.382 ·Ω^−1.683^, *V*2 (Ω) ≈ 0.621·Ω^−1.737^.

## Acknowledgements

The authors are grateful to Dr. James Quirk for help with implementing Bayesian programs for data analysis.

## Data and materials availability

Data is provided within the manuscript and also available upon request from the corresponding author.

